# Drones count wildlife more accurately and precisely than humans

**DOI:** 10.1101/165019

**Authors:** Jarrod C. Hodgson, Rowan Mott, Shane M. Baylis, Trung T. Pham, Simon Wotherspoon, Adam D. Kilpatrick, Ramesh Raja Segaran, Ian Reid, Aleks Terauds, Lian Pin Koh

## Abstract

Ecologists are increasingly using technology to improve the quality of data collected on wildlife, particularly for assessing the environmental impacts of human activities. Remotely Piloted Aircraft Systems (RPAS; commonly known as ‘drones’) are widely touted as a cost-effective way to collect high quality wildlife population data, however, the validity of these claims is unclear. Using life-sized seabird colonies containing a known number of replica birds, we show that RPAS-derived data are, on average, between 43% and 96% more accurate than data from the traditional ground-based collection method. We also demonstrate that counts from this remotely sensed imagery can be semi-automated with a high degree of accuracy. The increased accuracy and precision of RPAS-derived wildlife monitoring data provides greater statistical power to detect fine-scale population fluctuations allowing for more informed and proactive ecological management.

## Introduction

Wildlife populations are undergoing dramatic declines in response to a wide range of human-induced threats (Dirzo *et al.* 2014; Tilman *et al.* 2017). High quality ecological data are vital to monitor such changes. Emerging technologies, such as camera traps (Rowcliffe & Carbone 2008) and radio telemetry (Hussey *et al.* 2015; Kays *et al.* 2015), have increasingly been used to address this challenge (Moll *et al.* 2007; Hebblewhite & Haydon 2010; Pimm *et al.* 2015), especially when wildlife populations are rapidly fluctuating, highly mobile or located in remote habitats.

Remotely Piloted Aircraft Systems have been heralded as a game changer in ecology (Jones, Pearlstine & Percival 2006; Watts *et al.* 2010; Koh & Wich 2012; Anderson & Gaston 2013; Marris 2013; Chabot & Bird 2015; Linchant *et al.* 2015; Christie *et al.* 2016). They are used for data collection in an increasingly diverse suite of ecological applications, including identification of floristic biodiversity of understorey vegetation (Getzin, Wiegand & Schöning 2012), monitoring for poaching activities (Mulero-Pazmany *et al.* 2014), and bird surveys (Sarda-Palomera *et al.* 2012; Chabot, Craik & Bird 2015). However, there has been little consideration of the quality of data obtained using RPAS compared to more conventional methods (see Hodgson *et al.* (2016a) for an exception).

We assessed the accuracy of RPAS-facilitated wildlife population monitoring in comparison with the traditional ground-based counting method. The task for both approaches was to derive an estimate of the size (i.e. number of individuals) of 10 replica seabird colonies. Each replica colony had a different known number of life-sized individuals. We hypothesised that RPAS-derived counts would be more accurate and more precise than those generated using the traditional approach, confirming RPAS-technology as revolutionary for ecological monitoring.

## Materials and methods

### Study site and simulated colony set-up

Fieldwork (#epicduckchallenge) was completed at a metropolitan beach in South Australia (Port Willunga, 35°15’33 S, 138°27’41 E) in accordance with relevant permits (Department of Environment, Water and Natural Resources scientific research permit: M26523-1; City of Onkaparinga location permit: 4138). The experimental design, including the majority of anticipated statistical analyses, was pre-registered (Hodgson *et al.* 2016b).

Ten simulated Greater Crested Tern *Thalasseus bergii* breeding colonies were constructed using commercial, life-size, plastic duck decoys (∼ 25.5 x 11.3 cm, 185 cm^2^ footprint). Decoys provided a realistic representation of the nesting seabird stimuli observers encounter in the field. Colonies were situated separately on the beach, above the high water mark, in sandy areas that represented analagous nesting habitat. These were typically devoid of vegetation but often contained natural beach debris.

As inter-indiviudal interactions are thought to influence colony layout, a model of nesting pressure was applied to an underlying hexagonal grid to generate unique, unbiased colony layouts (Hodgson *et al.* 2016b). The hexagonal grid was re-created in the field using a wire mesh, upon which grid cell centres were marked (mean density: 11.39 m^-2^). Pre-counted wooden skewers were placed one per cell at a random location within all cells identified as occupied in the colony layout map. The mesh was removed and each skewer was replaced with a decoy facing approximately into the wind. The number of skewers retrieved was taken to be the true number of individuals in the colony. Colony sizes were between 463 and 1017 individuals. One individual was placed in each occupied cell.

### Ground counting approach

Ground counts were made by experienced seabird counters using a standard field technique (Hodgson *et al.* 2016a). Counters used tripod-mounted spotting scopes or binoculars as required. Hand-held tally counters were used to assist counting. Observation viewpoints at similar altitude to the colony and which provided the optimum vantage were selected (Fig. 1). Viewpoints were positioned 37.5 m from the nearest bird in the colony – this is the flight initiation distance for Caspian Tern *Hydroprogne caspia* (Moller *et al.* 2014) and so is a biologically-plausible minimum approach distance for a similar species in the field. Counts (*n* = 61) were 7 ± 2.65 min (s.d.) in duration. Four to seven counters each made a single blind count of the number of individuals in each colony. The numbers of counters were selected based on a preliminary power analysis (Hodgson *et al.* 2016b) which investigated the sample sizes necessary to detect small (∼ 10%) differences in mean counts and count variances between ground and RPAS-derived counts to high (80%, 90%, and 95%) power. Counters had no knowledge of the true number of individuals in colonies or the colony set-up technique. Counts were made between 0930 and 1645 on one day, resulting in variation in illumination and shadows.

**Figure 1:**
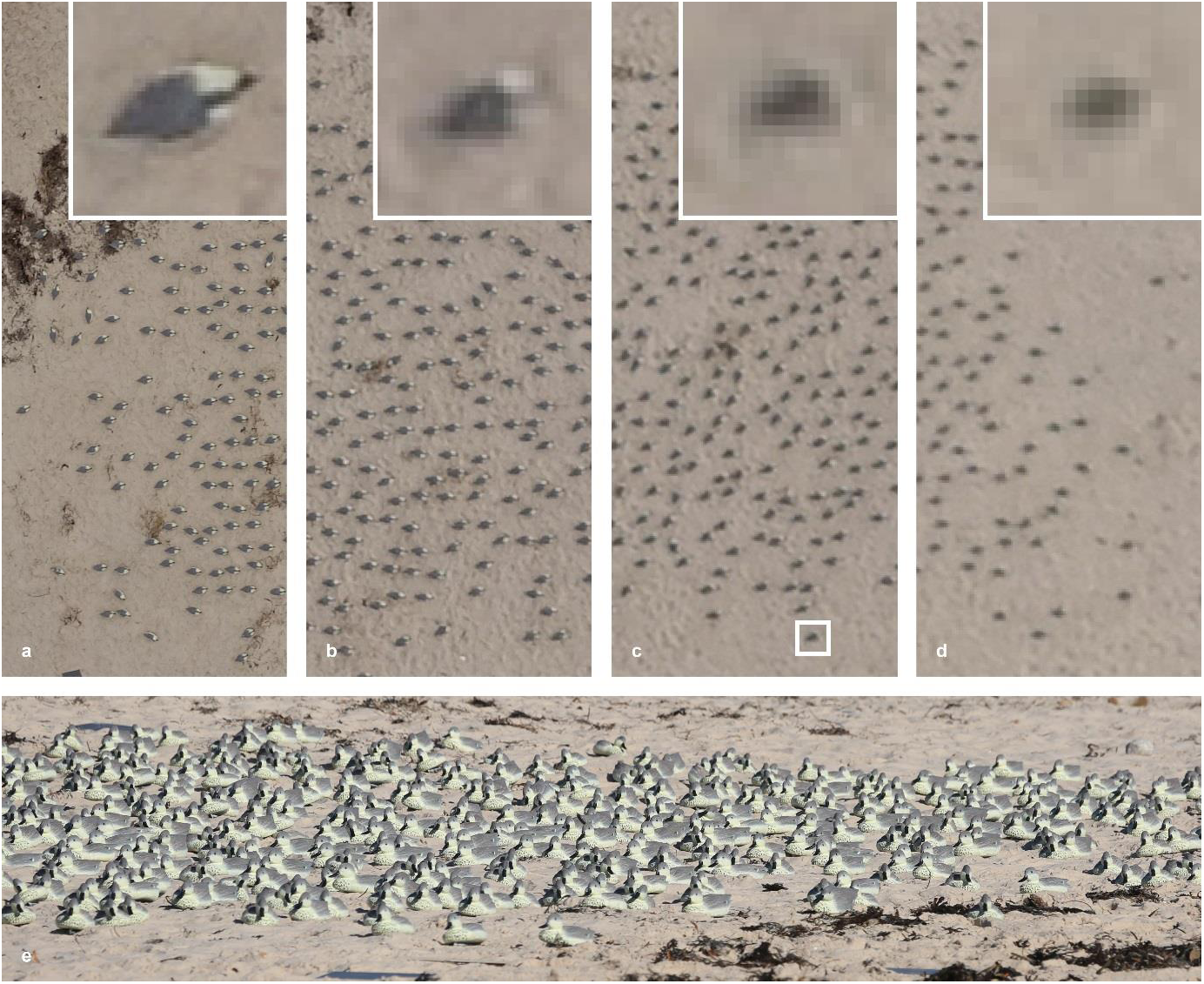
Aerial vantage of a replica seabird colony compared with the ground counter’s viewpoint. One colony represented by a mosaic of images (**a-d**) photographed from a RPAS-mounted camera at varying heights (30 m, 60 m, 90 m and 120 m) and resulting ground sample distances (GSD; 0.82 cm, 1.64 cm, 2.47 cm and 3.29 cm). Insets are of the same individual (square; **c**) at each height, displaying the decrease in resolution relative to an increase in GSD. **e**, View of the colony from a ground counter’s standing position.

### RPAS description, flight characteristics and data collected by RPAS

A small, off-the-shelf quadcopter (Iris+, 3D Robotics) was used as a platform to image each colony. After positioning the RPAS in the centre of the colony at 15 m above ground level, it was piloted in altitude hold mode to make a vertical ascent without movement in other axes. The RPAS was loitered for short periods (∼ 10 seconds) to enable the capture of several photographs at 30 m, 60 m, 90 m and 120 m above ground level (sample heights). Sampling was restricted to a height of 120 m as this is a common maximum limit for standard RPAS flight. Ground control station connection (Mission Planner, planner.ardupilot.com) was utilised and total flight time for missions was 5-7 min. All missions were in accordance with local regulations and flown by the same licenced pilot. Samples were collected within 40 min of the completion of ground counts.

Imagery was captured using a compact digital camera (Cyber-shot RX100 III, Sony – resolution: 5,472 x 3,648 px; sensor: CMOS; sensor size: 13.2 x 8.8 mm; lens: ZEISS Vario-Sonnar T). Exposure time was set at 1/2000 seconds using shutter priority mode. Photographs were captured successively (∼ 1 sec intervalometer) using the Sony PlayMemories Time-lapse application in jpeg format and at minimum focal length (8.8 mm). The camera was mounted facing downward using a custom vibration dampening plate. The footprint of a single image at each height encompassed the colony for all replicates. For analysis, only the image captured closest to the middle of the loiter time period for each sample height was used. These images (scenes; *n* = 40) were cropped (colony area < 50% of footprint) so that the image footprint was identical for each sample height for a given colony. High quality imagery was obtained for six of the ten colonies. Imagery for the remaining four colonies was affected by vibration-blur caused by a failure of the sensor attachment, likely due to wind speeds near the limit of the capability of the RPAS platform. Scenes are archived online (Pham & Hodgson 2017).

The ground sample distance (GSD), being the distance between adjacent pixel centres on the ground, for sample heights were 0.82 cm, 1.64 cm, 2.47 cm and 3.29 cm (Fig. 1). When photographed at nadir, this approximated to 275, 69, 30 and 17 pixels per individual respectively. The variance in GSDs was intended to represent the resolutions commonly achieved in wildlife monitoring applications, which result from sensor and sampling height variations.

### Manual RPAS image counting approach

Manual counts of perceived individuals in digital imagery were completed following a technique previously implemented for RPAS-derived monitoring of living seabirds (Hodgson *et al.* 2016a). Systematic counts were made using the multi-count tool within an open source, java-based scientific image processing program (ImageJ, <u>http://imagej.net/</u>). A grid plugin was used to overlay a square matrix (cell sizes: 70,000, 15,000, 8,000 and 4,000 pixels for each sample height) and counters were instructed to view the colony sequentially (gridcell-by-gridcell: left to right, top to bottom). Counters were encouraged to zoom in to each cell as they progressed and, upon completion, review their count at different levels of zoom until they were satisfied they had counted all individuals. For each sample height, seven to nine individuals counted each colony. Counters had no knowledge of the experimental setup and only one had experience ground counting colonial birds.

### Semi-automated aerial image counting approach

In each scene, digital bounding boxes were used to manually delimit a percentage of individual birds (Supplementary Fig. 1a). Areas of background were also delimited. These data were used to train a linear support-vector machine (a discriminative classifier; Cortes & Vapnik 1995), which predicted the likelihood of each pixel being a bird or background when applied to the corresponding scene (Supplementary Fig. 1b). Instead of relying on colour intensities, we computed rotation-invariant Fourier histogram of oriented gradient (Liu et al. 2013) features for each pixel used in the training processes. This resulted in the classifiers being trained to determine which features distinguished birds from the background. The predicted likelihood (score) maps indicated the approximate locations of birds in the scenes, and detections were generated by applying a threshold to the likelihood maps. This process unavoidably resulted in redundant bird proposals (Supplementary Fig. 1c) and so the final detection results were obtained by suppressing redundant proposals via minimising an energy function (Pham *et al.* 2016; Supplementary Fig. 1d). This function encoded the spatial distribution of objects and is informed by our knowledge of how the birds nest (e.g. two birds cannot occupy the same location). The source code and dataset are archived online (Pham & Hodgson 2017).

To determine the minimum amount of training data required for accurate detections relative to manual image counts, we varied the percentage of individual birds used as training data between 1% and 30% for each scene.

### Statistical methods

All analyses were carried out in *R* version 3.2.2 (R Core Team 2016). Pre-registered analyses were designed to investigate how within-colony absolute count error, within-colony variability of counts, and within-colony bias of counts differed between count techniques (Hodgson *et al.* 2016b).

For each test, a generalised linear mixed model was fit between the response (e.g. absolute count error) and the technology used to make the count (e.g. ground-count, manually counted RPAS at 30 m height, semi-automatically counted RPAS at 30 m height), with colony included in the model as a random effect (Supplementary Information 1). To investigate effects of counting technique on absolute count error, we defined the response as the absolute difference between the true number of birds in a colony and the counted number of birds. To investigate effects of counting technique on count-variability, we defined the response as the absolute difference between each count and the mean of counts of the same colony taken using the same method. Count variability was not estimated for semi-automated counts as there was only a single semi-automated count per colony. To investigate the effect of counting technique on relative count bias, we defined the response as the difference between the true number of birds in the colony and the counted number of birds. For the absolute count error model, we used a Quasipoisson distribution, and for the variability and bias models, we used a Gaussian distribution. For each model, post-hoc Tukey tests were used to test for differences in the response between all pairs of treatment levels.

Semi-automated count data were added to the experimental design subsequent to our pre-registration of the analysis which necessitated minor analysis modification. The addition of semi-automated count data, with a single replicate per colony, required fitting colony ID as a random effect instead of as a fixed effect in each model.

Statements comparing the accuracy of RPAS-derived counts to ground-based counts are based on the mean within-colony Root Mean Squared Error (RMSE) of that counting approach, standardised as a proportion of the true count within each colony (Supplementary Information 2). For instance, a statement that RPAS-derived counts are ‘95% more accurate than ground-counts’ means that, within-colony, the RMSE for RPAS-derived counts is 5% the RMSE for ground-based counts, representing a 95% reduction in RMSE.

To investigate the probability of counting each individual correctly, we developed models with a variety of possible counting outcomes for each object (Supplementary Information 3). We assumed that there are N = n0 + n1 + n2 objects that are counted by an observer, of which n1 are counted correctly, n0 are missed and n2 are double counted. We assumed that n = (n0, n1, n2) is multinomially distributed with probability p = (p0, p1, p2). In this structure, n0, n1, n2 are latent and the observer can only report the total count M = n1 + 2n2. Allowing each object to be at most double counted constrains n considerably, and the probability mass function can then be formulated. We adopted a Dirichlet prior for p (p∽Dirichlet(a)) making the conditional distribution of p: p|n(k) ∼ Dirichlet(a + n(k)).

We then ran a Gibbs sampling routine for p by alternately sampling n(k) and p from these two distributions. Since there was considerably more variation between ground counts compared to manual RPAS counts, we ran the analyses for each ground counter separately, and pooled data across counters for manual RPAS counts. The RPAS analyses were run separately for each sample height, and two sets of analyses were undertaken, one with data from all colonies, and the other with data from the subset of colonies with high quality imagery. This statistical approach does not account for objects being mistakenly identified as birds (i.e. false positives). Furthermore, as this model assumes each individual is counted independently, which may not always be true (particularly for ground counters who tend to count in clusters), care needs to be taken in the interpretation of the estimated probability values (and their variance).

To compare the semi-automated counts to that of the people counting the images, we first took the semi-automated count after 10% of training data had been used for each scene. Ten percent of training data was consistently identified as a threshold over which little improvement in counts occurred for all scenes. We compared this count to each of the manual counts of the same image using ANOVA for all scenes, and also for those scenes of high quality. We also used Poisson generalized linear models to make more quantitative comparisons of the two approaches.

### Code availability

R scripts used for analyses are available in the Supplementary Information.

### Data availability

The pre-registered experimental design is available via the Open Science Framework (*\*URL to be made public on publication\**). The count data, scenes and code for the image-analysis algorithm are archived online (*\*URL to be made public on publication\**).

## Results

### Manual RPAS-derived image counting versus ground counts

On average across all colonies, RPAS-derived counts were between 43% and 96% more accurate than ground counts, depending on the sample height (between 92% and 98% for the colonies with high quality imagery; Supplementary Table 1). The mean absolute error was significantly smaller for RPAS-derived counts at all heights compared to ground counts (all P < 0.001; Fig. 2a).

**Figure 2:**
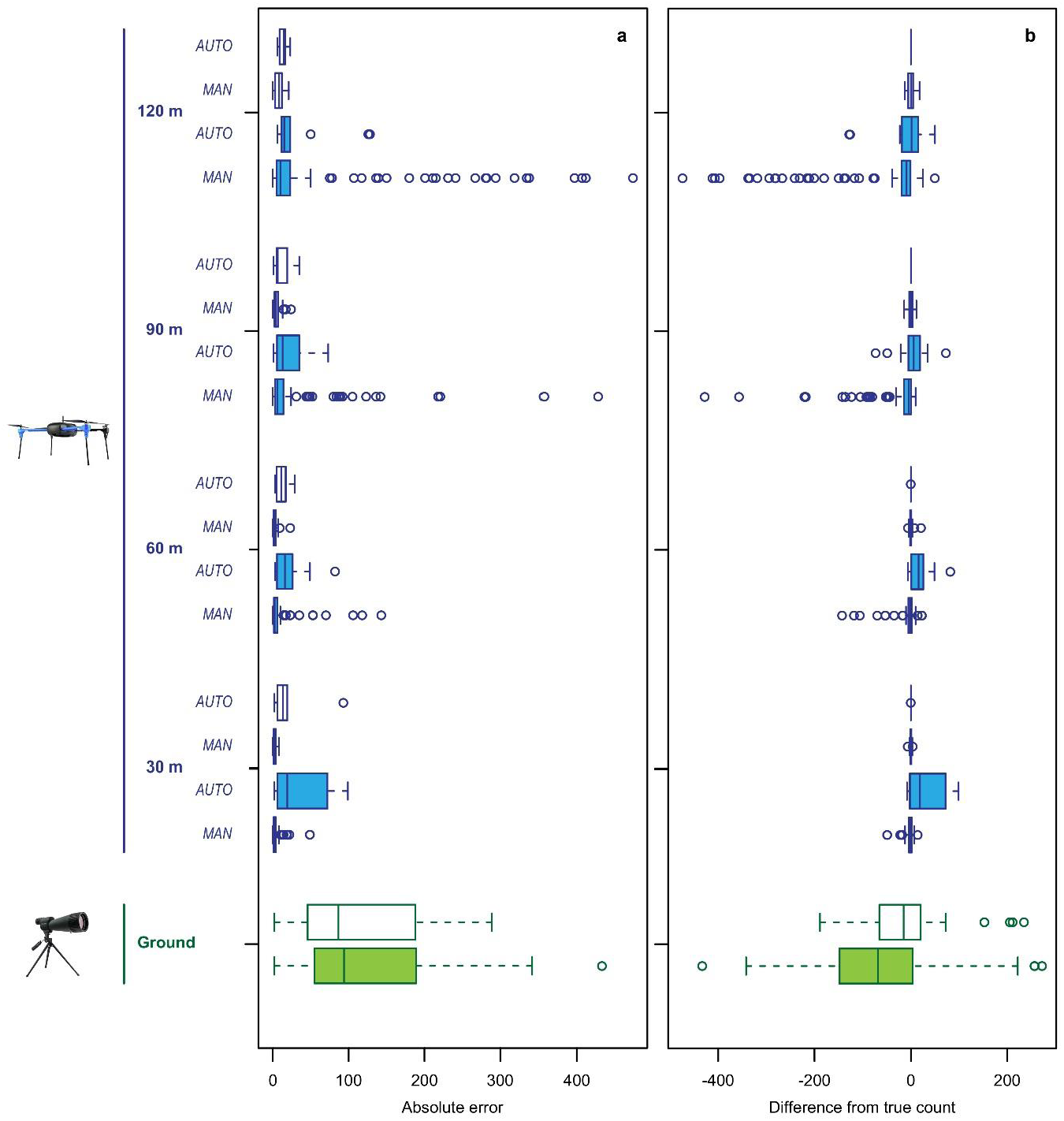
Accuracy and bias of RPAS and traditional wildlife monitoring approaches. The absolute error (**a**) and difference from the true count (**b**) of each method. Data from all colonies (*n* = 10; shaded) and also for the subset of colonies with high quality imagery (*n* = 6; unshaded) are presented for both RPAS-derived (blue) and ground (green) counts. RPAS-derived manual-human (Man) and semi-automated (Auto) counts are displayed and data are grouped by height which reflects ground sample distance (GSD; 30 m height = 0.82 cm GSD, 60 m = 1.64 cm, 90 m = 2.47 cm, 120 m = 3.29 cm).

No significant increase in count accuracy was achieved by obtaining imagery from heights lower than or equal to 90 m. Using data only from colonies with high quality imagery, there was no significant change in count accuracy across the range of heights. The lower accuracy of ground counts was due to significant underestimations of the true number of individuals in colonies (Fig. 2b). RPAS-derived counts from imagery obtained at 30 m and 60 m did not significantly under-or overestimate the true number of individuals in a colony, and there was no evident bias in RPAS-derived counts at any height for colonies with high quality imagery (Fig. 2b).

Using data from all colonies, RPAS-derived counts from 30 m and 60 m had a much higher probability (90% and 50%, respectively) of correctly counting an individual than counts from ground observers (< 10%) (Fig. 3). However, 90 m and 120 m probabilities were largely indistinguishable from the ground count probabilities, with a slightly higher likelihood of missing individuals compared to counting them twice (Supplementary Fig. 3, 4). Colony counts made from high quality imagery had a much higher probability of individuals being counted correctly, with > 85% probability of correctly counting an individual at all heights (Fig. 3). By contrast, ground counts had a low probability of counting an individual correctly, with the probability of double counting and missing an individual varying considerably between observers (Fig. 3 and Supplementary Fig. 3, 4).

**Figure 3:**
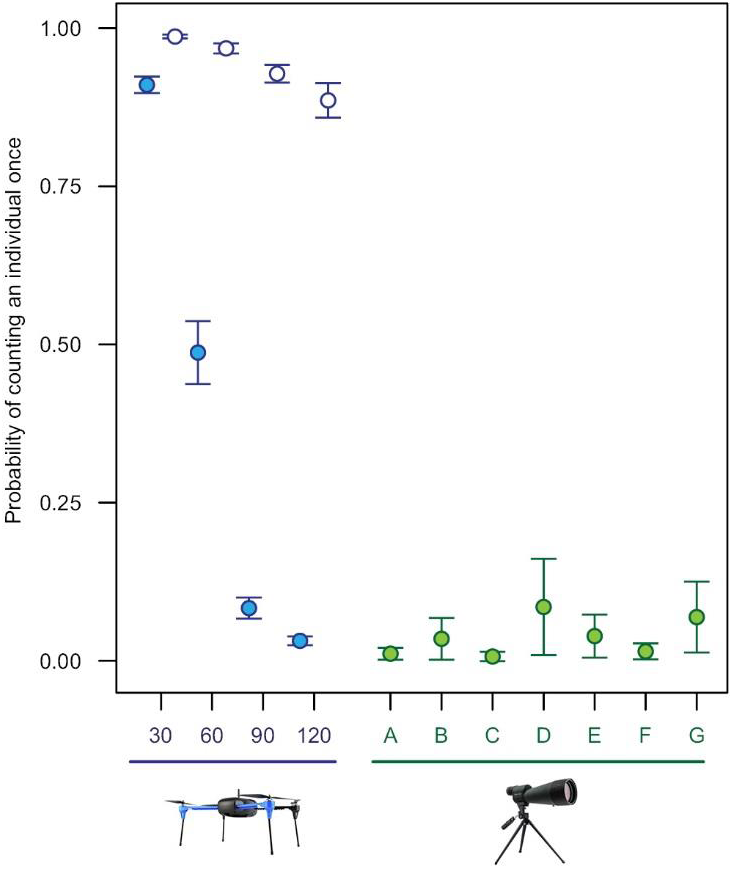
Probability of counting an individual in a colony once (correctly). Data from all colonies (*n* = 10; shaded) and also for the subset of colonies with high quality imagery (*n* = 6; unshaded) are presented for RPAS-derived (blue) manual counts. These data are grouped by height (m) which reflects ground sample distance (GSD; 30 m height = 0.82 cm GSD, 60 m = 1.64 cm, 90 m = 2.47 cm, 120 m = 3.29 cm). Probabilities from ground count data (green) for all colonies are estimated for each counter individually (A-G). Error bars represent standard deviation.

RPAS-derived counts were more precise (i.e. had lower inter-observer variability) than ground counts, regardless of the height at which imagery was obtained (t_4,560_-10.21 to-13.37, all P < 0.001; Supplementary Fig. 5). RPAS-derived counts were more precise for imagery obtained at 30 m compared to those obtained from 120 m (P = 0.01), however, there were no significant differences in precision among RPAS-derived counts at different heights for colonies with high quality imagery (all P > 0.98).

### Semi-automated RPAS approach

By increasing the percentage (1 – 30%) of individuals used as training data for the image-analysis algorithm, 10% training data was consistently identified as a threshold above which little improvement in count accuracy was achieved in this semi-automated approach (Supplementary Fig. 2). There was no significant difference between counts that were made with 10% training data and those made by manual counting of RPAS-imagery across all scenes. The semi-automated results were 94% similar to manual counts across all scenes (98% for the colonies with high quality imagery; see also Supplementary Table 1).

## Discussion

RPAS-derived data were more accurate and more precise than the traditional data collection method validating claims that RPAS will be a revolutionary tool for ecologists. Never has the importance of accurate wildlife population monitoring data been greater than at present given the alarming population declines observed in animal species across the globe (Dirzo *et al.* 2014). By facilitating accurate census, RPAS will provide ecologists with confidence in population estimates from which management decisions are made. Furthermore, the superior precision of RPAS-derived counts increases statistical power to detect population trends, owing to the lower type II error rate in statistical analysis that comes with comparing measures with smaller variance (Gerrodette 1987). The improved precision of wildlife population census using RPAS has been demonstrated for free-living seabird colonies (Hodgson *et al.* 2016a) suggesting our results are generalizable to natural settings. Differences in accuracy and precision between RPAS-facilitated and traditional survey methods can be attributed to the sources, and magnitude, of variance introduced into the two approaches which are strongly affected by the different vantages of the two methods (Hodgson *et al.* 2016a).

We have conducted two independent analyses of how count error differs across count approaches: a Frequentist analysis which estimates mean absolute count error, and a Bayesian analysis which estimates the probability of double-counting or missing individual animals. The two analyses are in agreement on the broad patterns: RPAS-derived counts are estimated to have lower error than ground counts in both analyses, and the error-rate is fairly insensitive to sample height for RPAS-derived counts. The Bayesian analysis makes restrictive assumptions about the process by which counting errors occur, and these assumptions may not fully reflect real-world counting processes. Nevertheless, the Bayesian analyses provide a first estimate of the extent to which overall count accuracy is dependent on the double-counting of some individuals cancelling out the effect of missing others. As an extreme example, our analysis suggests that < 10% of animals are correctly classified as a single animal in typical ground counts.

Manual counting of RPAS-derived imagery returned high quality data, but also involved substantial labour investments. Recent advances in digital sensors and image-analysis techniques have been increasingly employed to streamline the detection process (Chabot & Francis 2016). By applying a semi-automated image-based object detection algorithm to each scene, we vastly improved efficiency compared to the manual RPAS-derived census. Importantly, the reduction in person-hours provided by this semi-automated approach did not diminish data quality. This will be of particular interest in today’s research environment where funding for conservation is limited (Waldron *et al.* 2013) and researchers are under ever more pressing time commitments (Fischer, Ritchie & Hanspach 2012).

The capture quality and resolution of RPAS-derived imagery heavily influenced the results of both human and semi-automated detection. Consequently, ecologists should determine the minimum required GSD for their context, and optimise their sensor accordingly (e.g. resolution, focal length) relative to sample height. When determining an appropriate sample height, best practice protocols should be considered to minimise potential disturbance to wildlife (Hodgson & Koh 2016), while complying with relevant local aviation legislation and achieving an acceptable sample area within the possible survey time period.

The ability to collect data with higher accuracy, higher precision, and less bias than the existing approach confirms that RPAS are a scientifically rigorous data collection tool for wildlife population monitoring. This approach also produces a permanent record, providing the unique opportunity to error-check, and even recount with new detection methods, unlike ground count data. As RPAS platforms, sensors and computer vision techniques continue to develop, it is likely that the accuracy and cost effectiveness of RPAS-based approaches will also continue to improve.

## Acknowledgements

We particularly thank those who assisted with this study including Po-yun Wong, the ground support team and the numerous volunteers who made ground and RPAS-derived colony counts. We acknowledge the South Australian Department of Environment, Water and Natural Resources for granting a scientific research permit for this work. J.C.H. is supported by an Australian Government Research Training Program Scholarship. L.P.K. is supported by the Australian Research Council & Conservation International.

## Author contributions

J.C.H., R.M., S.M.B., A.T. and L.P.K. designed the study, analysed the data and wrote the manuscript. A.D.K. assisted with designing the study. J.C.H., R.M., S.M.B., A.D.K., R.R.S. and L.P.K. collected the data. T.T.P., J.C.H., L.P.K. and I.R. developed the semi-automated detection technique. S.W. and A.T. completed the multinomial analyses. All authors contributed to drafting the manuscript.

---

**Supplementary Figure 1:**
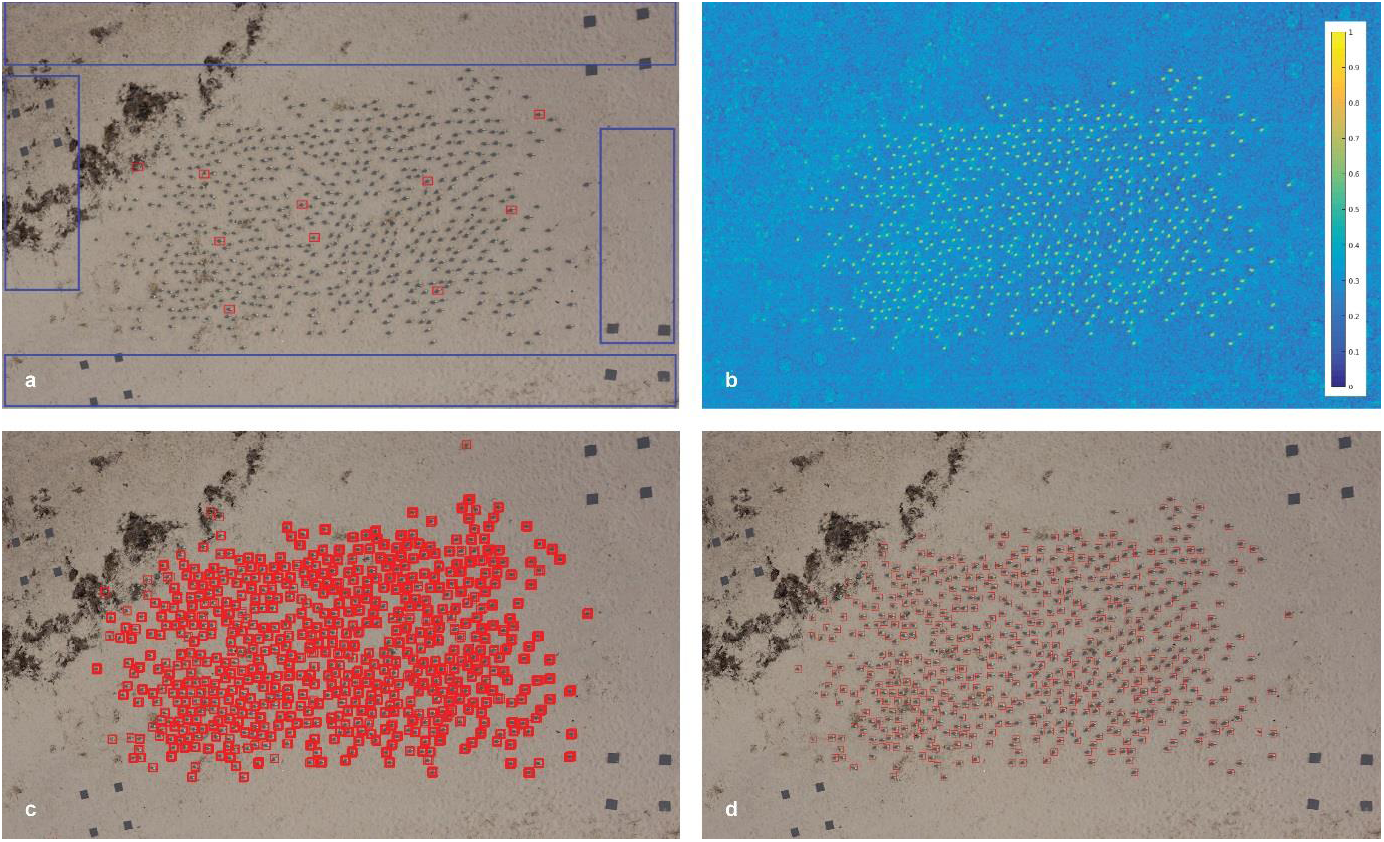
Semi-automated detection and counting of wildlife using computer vision techniques. (**a**) User annotation of perceived target objects (red) and background (blue). (**b**) score map generated by the trained classifier which has automatically determined which image features distinguish objects from background, independent of scale and orientation. Warmer colours indicate increasing likelihood of the pixel being a target object. (**c**) target object proposals (red) computed by thresholding the score map. Object size is estimated from the annotations. (**d**) final output (which includes a total count and detection co-ordinates) where detected individuals are delineated (red) after redundant detections have been automatically suppressed.

**Supplementary Figure 2:**
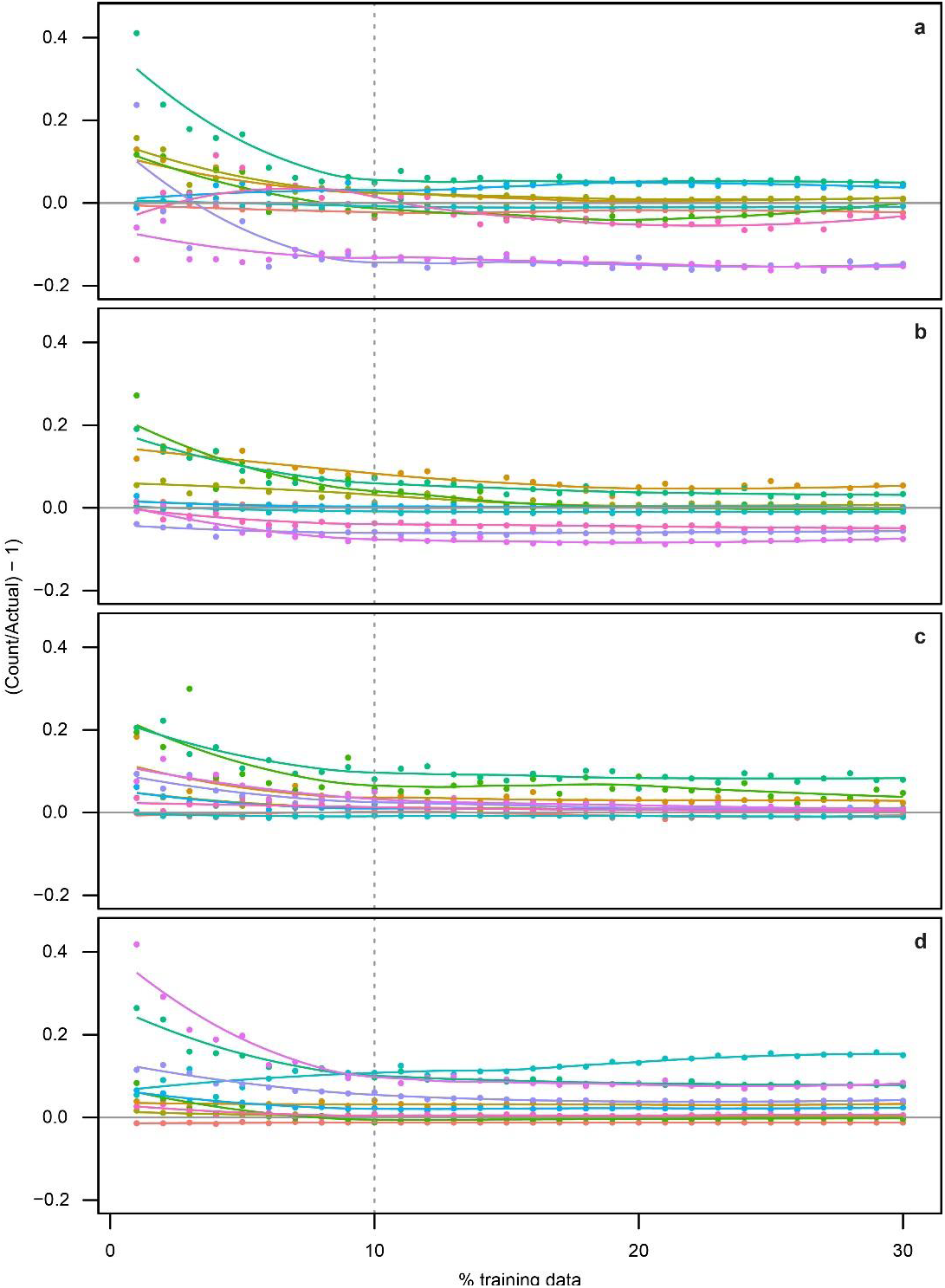
Improvement in accuracy of semi-automated detection counts with increasing training data. Colonies (*n* = 10) are represented by individual colours at each height which reflects ground sample distance (GSD): 120 m height = 3.29 cm GSD (**a**); 90 m = 2.47 cm (**b**); 60 m = 1.64 cm (**c**); 30 m = 0.82 cm (**d**). Lowess smoothed trendlines are displayed. Analyses were computed using count estimates generated from 10 % training data (dashed line).

**Supplementary Figure 3:**
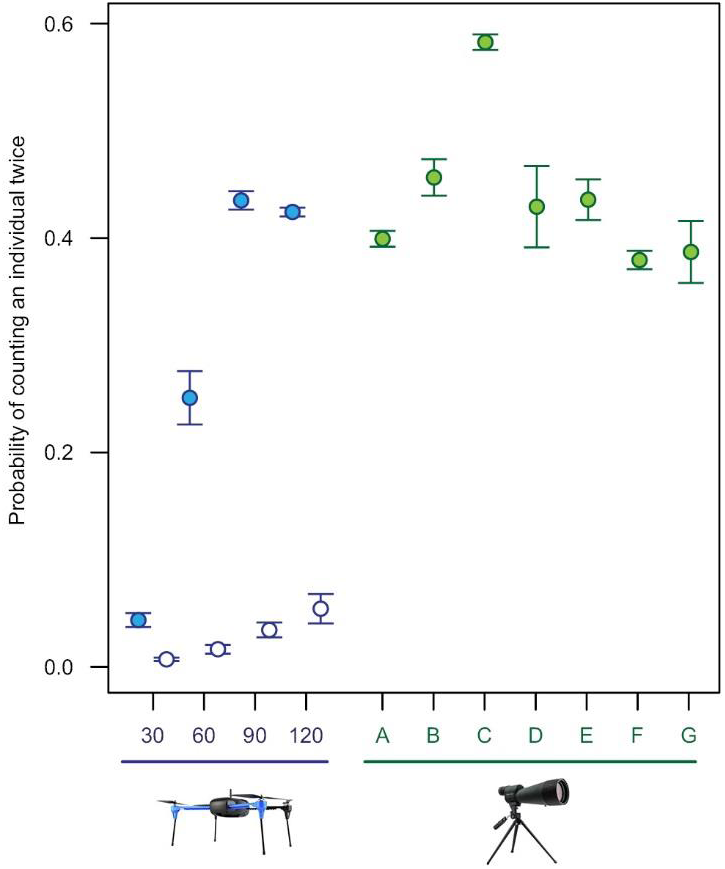
Probability of counting an individual in a colony twice (double counting). Data from all colonies (*n* = 10; shaded) and also for the subset of colonies with high quality imagery (*n* = 6; unshaded) are presented for RPAS-derived (blue) manual counts. These data are grouped by height (m) which reflects ground sample distance (GSD; 30 m height = 0.82 cm GSD, 60 m = 1.64 cm, 90 m = 2.47 cm, 120 m = 3.29 cm). Probabilities from ground count data (green) for all colonies are estimated for each counter individually (A-G). Error bars represent standard deviation.

**Supplementary Figure 4:**
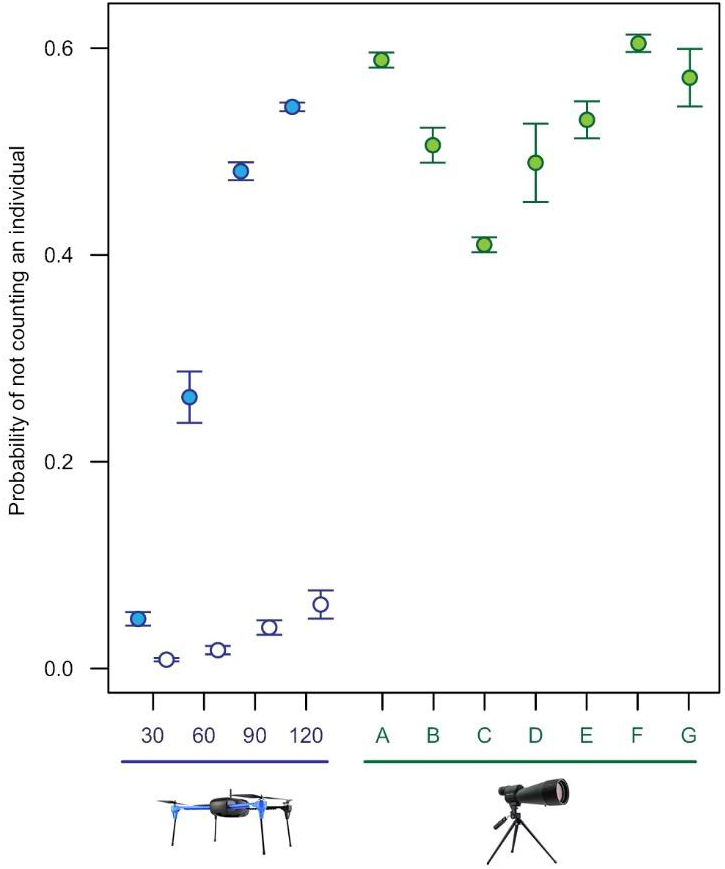
Probability of not counting (missing) an individual in a colony. Data from all colonies (*n* = 10; shaded) and also for the subset of colonies with high quality imagery (*n* = 6; unshaded) are presented for RPAS-derived (blue) manual counts. These data are grouped by height (m) which reflects ground sample distance (GSD; 30 m height = 0.82 cm GSD, 60 m = 1.64 cm, 90 m = 2.47 cm, 120 m = 3.29 cm). Probabilities from ground count data (green) for all colonies are estimated for each counter individually (A-G). Error bars represent standard deviation.

**Supplementary Figure 5:**
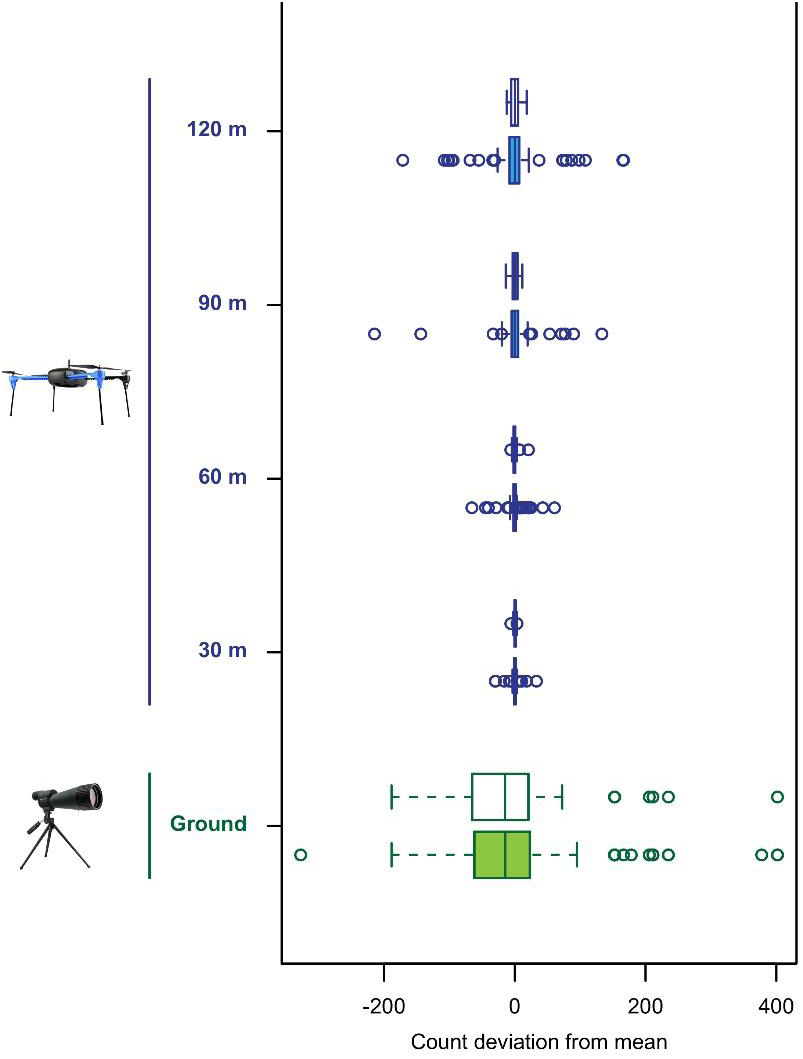
Precision of RPAS and traditional wildlife monitoring approaches. Data from all colonies (*n* = 10; shaded, lower box in each course) and also for the subset of colonies with high quality imagery (*n* = 6; unshaded, upper box in each course) are presented for both RPAS-derived manual-human (blue) and ground (green) counts. Data are grouped by height which reflects ground sample distance (GSD; 30 m height = 0.82 cm GSD, 60 m = 1.64 cm, 90 m = 2.47 cm, 120 m = 3.29 cm).

**Supplementary Table 1:**
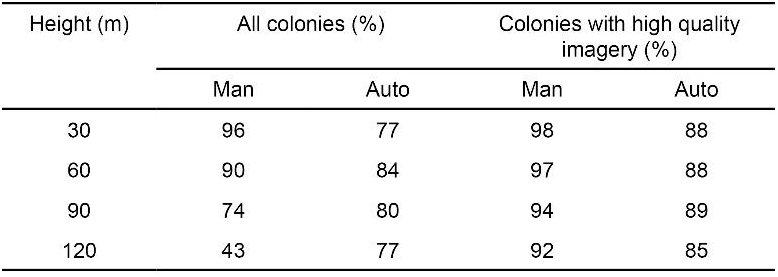
Mean percentage increase in accuracy of RPAS wildlife monitoring approaches compared with the traditional ground count approach. Percentages are calculated for RPAS-derived human manual (Man) and semi-automated (Auto) counts using data from all colonies (*n* = 10) as well as the subset of colonies with high quality imagery (*n* = 6). Data are grouped by height which reflects ground sample distance (GSD; 30 m height = 0.82 cm GSD, 60 m = 1.64 cm, 90 m = 2.47 cm, 120 m = 3.29 cm).

